# The nucleotide analog bemnifosbuvir inhibits hepatitis E virus replication in preclinical models

**DOI:** 10.1101/2025.08.15.670060

**Authors:** Jungen Hu, Tianxu Liu, Mara Klöhn, Andrew Freistaedter, Elif Toprak, Huanting Chi, Paula Jordan, Xinyue Yang, Johanna Becker, Volker Lohmann, Eike Steinmann, Lin Wang, Viet Loan Dao Thi

## Abstract

**Background:** Hepatitis E virus infections remain a global health concern. Immunocompromised patients are at an increased risk to develop chronic HEV infection and thereby severe liver disease. Current off-label regimens are suboptimal with treatment failure being reported. Therefore, there is an urgent need for an effective anti-HEV treatment.

**Objective:** In this study, we aimed to identify potent inhibitors of HEV replication.

**Design:** We developed a rapid, image-based screening platform based on a full-length HEV fluorescence reporter virus and screened a nucleotide/nucleoside analog library. The identified lead candidate was validated in authentic hepatocyte culture systems, as well as in a gerbil infection model.

**Results:** Bemnifosbuvir (BEM), previously characterized as a nucleotide analog with activity against other RNA viruses, efficiently suppressed HEV replication *in vitro* and in *vivo* in a dose-dependent manner, with minimal cytotoxicity at effective concentrations. Combining BEM with ribavirin, the off-label drug given to chronic HEV patients, resulted in an additive antiviral effect against HEV. We found that HEV-3 remains susceptible to inhibition by BEM over an extended treatment period, reducing concerns about the rapid development of viral resistance. Importantly, BEM significantly reduced HEV viral loads and liver inflammation in a gerbil infection model.

**Conclusions:** Given BEM’s favorable safety profile in preclinical and clinical settings, our results suggest investigating its efficacy in patients with chronic HEV infection.

## Introduction

Annually, hepatitis E virus (HEV) infects approximately 20 million people worldwide and is one of the most common causes of acute viral hepatitis^1^. Belonging to the *Hepeviridae* family, HEV can infect a wide range of animal species, with humans primarily infected by genotypes 1 through 4 (HEV-1 to 4) from the species *Paslahepevirus balayani*. HEV-1 and HEV-2 are transmitted primarily fecal orally between humans. In contrast, most HEV-3 and HEV-4 infections have zoonotic origins, often acquired through the consumption of contaminated meat products^2^. Consequently, HEV-1 and -2 are more prevalent in developing countries, while HEV-3 and -4 infections are more common in developed countries.

In healthy individuals, all four genotypes typically cause asymptomatic, self-limiting infections^2^, with fulminant hepatitis being rare^3^. However, HEV-3 and HEV-4 have been associated with chronic infection—defined as persistent viremia lasting more than 3–6 months—in immunocompromised populations^4 5^. These patients are at increased risk of rapid liver disease progression, including fibrosis and cirrhosis^5^.

Despite growing awareness and the pressing medical need, a direct-acting antiviral specifically targeting HEV has yet to be developed. Currently, chronic HEV infections are managed off-label with ribavirin (RBV) and pegylated interferon-α (IFN-α), with variable success rates^6^. However, IFN-α therapy can cause significant side effects and is generally contraindicated in transplant recipients due to the risk of graft rejection^6^.

Nucleoside analogues have revolutionized the treatment of various chronic viral infections by mimicking natural nucleosides, leading to chain termination or defective viral replication when incorporated into the viral genome^7^. Their structural similarity to natural nucleosides makes them promising candidates for broad-spectrum antiviral development. However, RBV achieves sustained virological responses in only about 80% of chronic HEV cases and is often associated with severe adverse effects^6^. Moreover, mutations—particularly in the viral RNA-dependent RNA polymerase (RdRp) gene—have been linked to therapy failure^8 9^. Likewise, an amino acid substitution in the HEV genome was reported to confer resistance against hepatitis C virus (HCV) polymerase inhibitor sofosbuvir (SOF)^10^, which we previously described to inhibit HEV replication *in vitro*^11^. Therefore, there is an urgent need for the development of a potent and specific anti-HEV treatment with a high resistance barrier.

The ~7.2 kb positive sense HEV RNA genome encodes three major open reading frames (ORFs): ORF1, the viral replicase; ORF2, the capsid protein; and ORF3, a small phosphoprotein essential for viral particle release^12^. High-throughput screens (HTS) to identify HEV inhibitors have traditionally relied on sub-genomic replicons, where regions encoding ORF2 and ORF3 are replaced with reporter genes such as GFP or Gaussia luciferase^13 14^. However, recent findings indicate that the structural proteins, particularly ORF2, also play critical roles in counteracting cell-intrinsic antiviral responses, thereby facilitating authentic viral replication^15^.

To address these limitations, we engineered a novel full-length HEV-3 fluorescence reporter virus by inserting a small GFP-11 tag^16^ at the C-terminus of the ORF2 capsid protein. Using this reporter system, we screened a library of nucleoside analogues and identified bemnifosbuvir (BEM), a drug that has shown potent pan-genotypic activity against HCV both preclinically and clinically^17 18^, as our top hit. Our study demonstrates that BEM potently inhibits HEV replication both *in vitro* and *in vivo*, using a gerbil infection model^19^. Remarkably, HEV replication remained sensitive to the treatment even after 40 serial passages and spanning over 160 days in cell culture with intermittent exposure to BEM. These results mitigate concerns about the development of viral resistance and underscore the potential of BEM as a promising treatment for chronic HEV infections.

## Materials and methods

### Reagents and plasmids

The nucleotide compound library (HY-L044) was purchased from MedChemExpress as pre-dissolved solutions in DMSO. BEM (T9341, TargetMol), NITD008 (HY-12957, MedChemExpress), SOF (HY-15005, MedChemExpress) and RBV (HY-B0434, MedChemExpress) were dissolved in DMSO. Interferon-α 2a (IFNα2a) (11100, PBL Assay Science) was dissolved in PBS containing 0.1% bovine serum albumin. All compounds were stored at – 80 °C.

HEV-3 ORF2::GFP11 and its replication-deficient GNN mutant were generated based on the HEV-3 Kernow-C1 p6 strain (a kind gift from Suzanne Emerson, GenBank accession number: JQ679013.1). The insertion and mutation were introduced by overlap extension PCR with Phusion polymerase (New England Biolabs) using primers listed in Supplementary table 1 or by restriction digest and ligation from available mutants^15^.

**Table 1:** Top 6 hits identified in the screen.

| No. | Name | Normalized inhibition (%) | Normalized cell viability (%) |
| --- | --- | --- | --- |
| 1 | Bemnifosbuvir | 97.67 | 94.40 |
| 2 | 3-Deazaadenosine hydrochlorid | 74.58 | 99.52 |
| 3 | 5'-O-TBDMS-dT | 66.20 | 93.68 |
| 4 | Sofosbuvir | 65.64 | 103.83 |
| 5 | MK-0608 | 59.76 | 103.64 |
| 6 | Uridine triphosphate trisodium salt | 57.86 | 103.11 |

### Nucleotide compound library screening

S10-3 GFP1-10 cells were electroporated with HEV ORF2::GFP11 RNA and seeded in cell culture flasks. The next day, 3×10^3^ cells were re-seeded into 96-well plates (3904, Corning). 48 h post-electroporation, the prediluted library compounds were added to the cells at a final drug concentration of 10 μM (exceptionally due to lower stock concentrations, enocitabine, ribavirin carboxylic acid, and cyclic AMP were at 2 μM; ATP disodium salt hydrate was at 3 μg/mL) in 0.5% DMSO. 5 μM NITD008 was added as positive control and 0.5% DMSO as negative control. 72 h later, the cells were fixed with 2% PFA for 10 min at room temperature (RT) and stained with Hoechst 33342 (1:2000 diluted in PBS). The cells were washed three times with PBS and then imaged using the Tecan Spark Cyto imaging plate reader with a 10x objective. Nuclei were identified by Hoechst staining, and GFP signals were analyzed using the Tecan ImageAnalyzer software. Briefly, primary masking of nuclei was performed based on the blue channel, with a sensitivity set to 70% and size parameters of 8– 30 μm for length and width. Secondary masking was conducted on the green channel, using a radius of 30 μm from the center of each nucleus, Voronoi masking, and a threshold of 0.01 relative fluorescence units. Normalized inhibition and cell viability were calculated based on the NITD008 and DMSO controls, respectively.

### HEV infection and drug inhibition assays

The cells were seeded, and the following day, inoculated with HEV in Minimum Essential Medium (MEM) supplemented with 10% FBS for the S10-3 cells or hepatocyte culture medium (HCM, Lonza) for the HLCs, and left to incubate overnight. The inoculum was removed, and the infected S10-3 cells and HLCs were treated with drugs. The medium was replaced every other day.

For serial passaging, the infected S10-3 cells were treated with different concentrations of BEM (0.4, 2, and 10 μM) starting on day 7 post-infection. Every four days, the cells were split 1:2 and passaged into a new well. The cells were treated alternating with or without drugs every three passages. PHHs were inoculated with HEV and drugs in HHMM for three days. Electroporated S10-3 GFP1-10 cells were treated with drugs for three days starting on day two post-electroporation.

### Gerbil infection model

13-week-old adult Mongolian gerbils (*Meriones unguiculatus*) (3 males and 3 females) were randomly selected and purchased from Sipeifu Biotech (Beijing, China). Serum and fecal samples were collected weekly and tested for the presence of HEV RNA and/or anti-HEV antibodies to exclude current or previous HEV infections before use in our study. The animals were housed individually in separate cages with adequate supplies of food and water.

HEV copy numbers were quantified in the HEV-positive inoculum by RT-qPCR. The negative inoculum was prepared from the feces of an uninfected gerbil. Each gerbil was injected with 1 mL of the inoculum intraperitoneally (i.p.). The daily dosage of BEM was dissolved in DMSO and diluted with water to 500 μL. It was administered at 250 mg/kg/d via gavage once daily for ten days, beginning on day five post-inoculation. The same volume of solvent was used as a vehicle control.

Fecal samples were collected every 3 to 4 days and stored at −80°C. Tissue specimens were collected when the gerbils were sacrificed. No animal was excluded from the analysis.

### Ethics approval statement

Use of human induced pluripotent stem cells (S-439/2018) and the HEV-3 patient isolate (S-292/2019) has been approved by the Ethics Committee of the Medical Faculty of Heidelberg University. For the PHHs, patient informed consent was obtained by Primacyt, as stated on their website. The gerbil experiment (LA2024168) has been approved by the Committee of Laboratory Animal Welfare and Ethics, Peking University Health Science Center.

### Patient and Public Involvement

Patients or the public were not involved in the design, or conduct, or reporting, or dissemination plans of our research.

### Data analysis and statistics

For the combination analysis, synergy scores are based on ZIP synergy analysis using SynergyFinder software programmed with R^20^. Graphs were made and statistical analyses were performed using GraphPad Prism 8. In all figures where p-values were calculated, the corresponding statistical test is listed in the figure legend.

For further details regarding the materials and methods used, please refer to the supplementary information.

## Results

### A novel full-length HEV fluorescent reporter virus can recapitulate the entire viral life cycle

Previous HTS studies have relied on HEV sub-genomic replicons, in which the regions encoding ORF2 and ORF3 have been replaced with reporter genes. Since these viral proteins are thought to play considerable roles in counteracting antiviral responses^15 21^, we aimed to engineer a full-length HEV reporter virus based on the cell culture adapted HEV-3 Kernow C1 p6 strain^22^ and a split GFP system^16^.

To this end, we took advantage of two observations. First, the ORF2 protein is highly expressed in HEV-infected cells and second, the ORF2 protein within the full-length genome can be tagged via duplication of the 3’ end^23^. However, since the duplication does not tolerate large inserts, such as full-length GFP, we tagged ORF2 with a small split GFP11 peptide (residues 215–230) through a flexible linker at its C-terminus (HEV-3 ORF2::GFP11, **Figure 1A**). In order to trans-complement the fluorescent molecule, we transduced hepatoma S10-3 cells with lentivirus to ectopically express the GFP1-10 fragment (residues 1–214, **Supplementary Figure 1A**).

**Figure 1.**
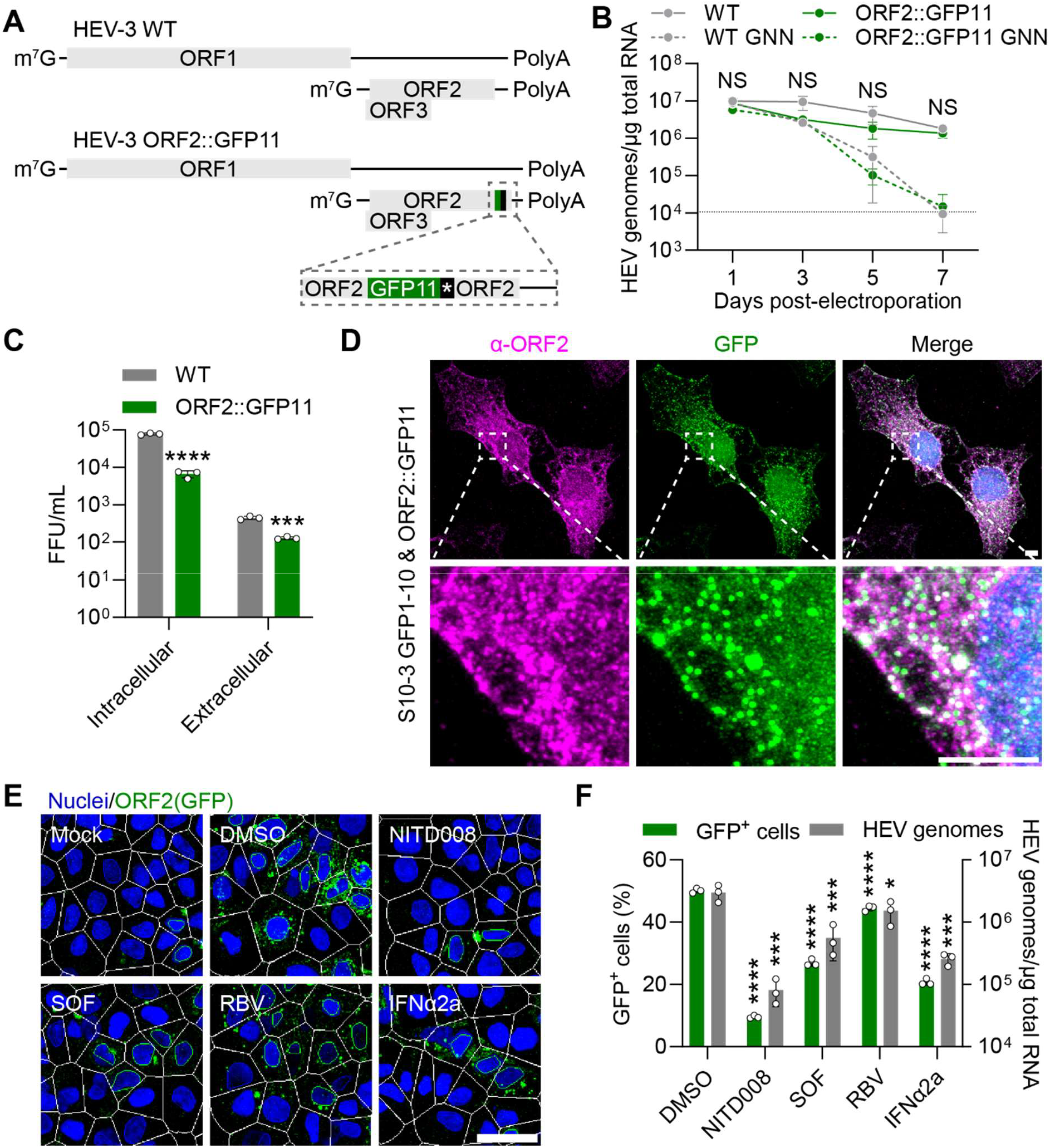
A novel full-length HEV fluorescence reporter virus can recapitulate the entire viral life cycle. (**A**) Schematic genome structure of HEV-3 WT (based on Kernow-C1 p6) and HEV-3 ORF2::GFP11. (**B**) S10-3 GFP1-10 cells were electroporated with *in vitro* transcribed HEV RNA and intracellular HEV genomes were quantified by RT-qPCR. GNN = a non-replicating mutant in the GDD motif. Data show mean ± SD of n = biological replicates from 3 independent experiments. Statistical significance was tested between WT and ORF2::GFP11 using unpaired Student’s t-test of each time point independently, adjusted for multiple testing using the Benjamini-Hochberg method. NS, non-significant. (**C**) Intracellular and extracellular virions were harvested from electroporated S10-3 cells 7 d post-electroporation and titered on HepG2/C3A cells. 7 d post-infection, HepG2/C3A cells were fixed and stained against ORF2 protein. Entire cell culture wells were imaged and foci forming units (FFU) were counted manually. Data show mean ± SD of n = biological replicates from 3 independent experiments. Statistical analysis was performed using unpaired two-tailed Student’s t-test. ***, p < 0.001; ****, p < 0.0001. (**D**) S10-3 GFP1-10 cells electroporated with *in vitro* transcribed ORF2::GFP11 RNA were fixed and stained against ORF2 protein (magenta) and nuclei (blue) 5 d post-electroporation. Shown are maximum projections of 40 confocal slices. Scale bar = 5 μm. (**E**) S10-3 GFP1-10 cells electroporated with ORF2::GFP11 RNA were treated with DMSO (0.5% v/v), NITD008 (5 μM), sofosbuvir (SOF, 10 μM), ribavirin (RBV, 10 μM) or interferon-α 2a (IFNα2a, 1000 IU/mL) 2 d post-electroporation. 3 d later, cells were fixed, stained with Hoechst 33342, and imaged with a Tecan Spark Cyto imaging plate reader. Segmentation and analysis were performed with Tecan ImageAnalyzer software. Shown are representative images. White outline = cell segmentation; Green outline = GFP^+^ cell; Blue outline = GFP^-^ cell. Scale bar = 50 μm. (**F**) Shown are % of GFP^+^ cells from (E) and HEV genomes quantified by RT-qPCR on day 5 post-electroporation. Data show mean ± SD of n = biological replicates from 3 independent experiments. Statistical analysis was performed using one-way ANOVA using Tukey’s test *post hoc*. *, p < 0.05; ***, p < 0.001; ****, p < 0.0001.

First, we compared the replication of HEV-3 ORF2::GFP11 with that of HEV-3 wild-type (WT) virus. As shown in **Figure 1B**, both viruses replicated to similar levels but not their replication deficient counterparts in which we mutated the critical “GDD” motif within the RdRp to “GNN”. We also titered infectious progeny collected from either the cell lysate (naked, intracellular HEV particles) or from the supernatant (quasi-enveloped, extracellular HEV particles) from S10-3 cells electroporated with HEV-3 ORF2::GFP11 or HEV-3 WT virus. Both intra- and extracellular HEV-3 ORF2::GFP11 particles were infectious, although at lower titers compared to HEV-3 WT particles (**Figure 1C**). These results, together with Western Blot analysis (**Supplementary Figure 1B**), showed that replication of the reporter virus led to the expression of all viral proteins. They further suggested that, although the GFP11 tag may impair some ORF2 functions, the reporter virus can still recapitulate the entire HEV life cycle.

When S10-3 cells expressing GFP1-10 were electroporated with HEV-3 ORF2::GFP11 RNA, the two fragments reassembled to produce a bright fluorescent ORF2 protein (**Figure 1D**), allowing for the convenient and rapid quantification of replicating cells. Next, we aimed to develop an inexpensive, rapid, image-based HTS platform to identify novel anti-HEV inhibitors. To this end, we used a Tecan Spark Cyto imaging plate reader with built-in segmentation, which enabled us to image and analyze a 96-well plate in less than 30 minutes. We used this reader to detect and analyze GFP^+^ cells as an indicator of HEV infection and nuclei as an indicator of cell viability (**Figure 1E**).

We first tested the sensitivity of HEV-3 ORF2::GFP11 infection to previously described HEV inhibitors through the quantification of GFP-positive cells. The tested inhibitors— including nucleoside analogs NITD008^24^, SOF, and RBV, as well as IFNα2a— all effectively suppressed HEV-3 ORF2::GFP11 replication, as evidenced by a similar reduction in number of both viral genomes and GFP^+^ cells (**Figure 1F**). These findings showed that our HEV-3 ORF2::GFP11 reporter virus-based platform was suitable for the screening of antiviral compounds.

### An image-based nucleoside/nucleotide library screen reveals bemnifosbuvir as a potent inhibitor of HEV replication

Next, we used the HEV-3 ORF2::GFP11 reporter to screen a commercial library of 539 nucleoside/nucleotide analogs. The adenosine analog NITD008 was previously reported to strongly inhibit HEV replication *in vitro*^24^. Although NITD008 exhibited toxicity in animal studies^25^ and was not evaluated further in clinical trials, we selected it as a positive control for our screening due to its potent *in vitro* activity. As shown in **Figure 2A**, S10-3 cells electroporated with HEV-3 ORF2::GFP11 RNA and treated with increasing concentrations of NITD008 exhibited a dose-dependent reduction in GFP^+^ cells (green line), without affecting the number of nuclei (gray line).

**Figure 2.**
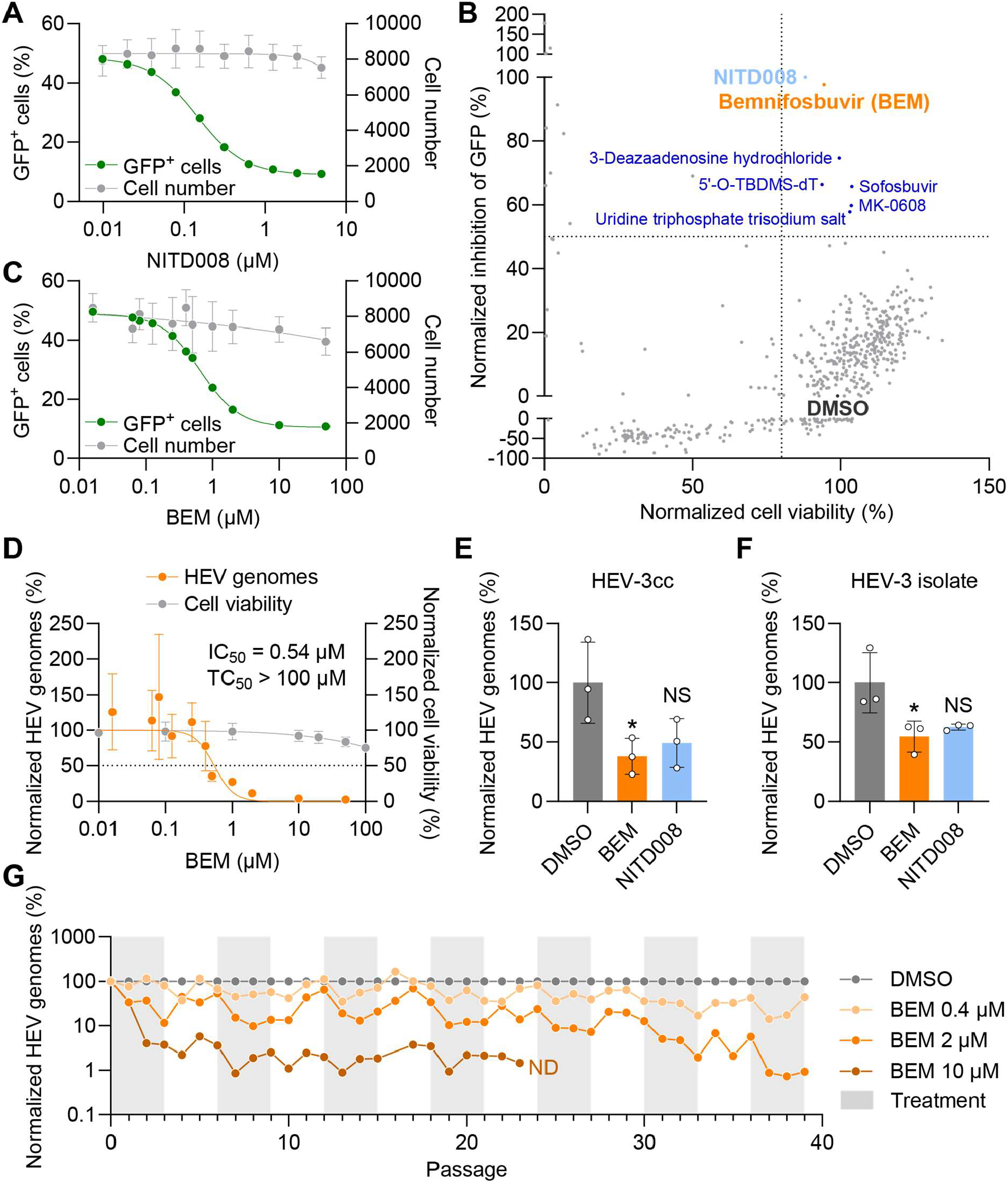
An image-based nucleoside/nucleotide library screen reveals bemnifosbuvir as a potent inhibitor of HEV replication. (**A**) S10-3 GFP1-10 cells electroporated with ORF2::GFP11 RNA were treated with different concentrations of NITD008 for 72 h from day 2 post-electroporation. 0.5% DMSO was used as vehicle control. The percentage of GFP^+^ cells was analyzed as described for Figure 1. Data show mean ± SD of n = biological replicates from 3 independent experiments. (**B**) Nucleoside/nucleotide library screening was performed using the same setup as described in (A). 0.5% DMSO and 5 μM NITD008 were used as negative and positive controls, respectively. (**C**) S10-3 GFP1-10 cells were treated with BEM as described in (A). Data show mean ± SD of n = biological replicates from 3 independent experiments. (**D**) S10-3 cells were infected with HEV-3cc WT virus (MOI=1 FFU/cell). The next day, viral inoculum was removed and cells were treated with increasing concentrations of BEM for 72 h. Intracellular HEV genome copies were quantified by RT-qPCR and cell viability was assessed. 0.5% DMSO was used as vehicle control. Data show mean ± SD of n = biological replicates from 3 independent experiments. (**E, F**) Induced pluripotent stem cell-derived hepatocyte-like cells were infected with (E) cell culture-derived HEV-3cc WT virus (MOI=1×10^6^ genome copies/well) or (F) a patient-derived HEV-3 isolate (MOI=3×10^2^ genome copies/well) and treated with 10 μM BEM or 5 μM NITD008 1 d post-infection for 144 h. Intracellular HEV genome copies were quantified by RT-qPCR. Data show mean ± SD of n = 3 biological replicates. Statistical analysis was performed using one-way ANOVA using Tukey’s test *post hoc*. *, p < 0.05; NS, non-significant. (**G**), S10-3 cells were infected with HEV-3cc WT virus (MOI = 1 FFU/cell) and intermittently treated for three consecutive passages with or without indicated concentrations of BEM. At each passage, cells were split 1:2 and HEV genome copies were harvested from lysates and analyzed by RT-qPCR. n = 1 biological replicate. ND, non-detected.

For the library screen, S10-3 cells were electroporated with *in vitro* transcribed HEV-3 ORF2::GFP11 RNA and seeded into 96-well microplates. After 48 h, 10 μM of each nucleoside/nucleotide analog was added to individual wells, with the exceptions previously noted. 3 days post-treatment, cells were fixed, imaged, and analyzed using the imaging plate reader. The number of GFP-positive cells served as a measure of antiviral activity, while the total nuclei count was used to assess drug-induced cytotoxicity.

As shown in **Figure 2B**, we identified six compounds that inhibited HEV replication without compromising cell viability (**Table 1**). Among these, bemnifosbuvir (BEM) emerged as the most promising candidate due to its strong capacity to suppress HEV replication, being close to that of the positive control NITD008. BEM has demonstrated potent activity against HCV across genotypes *in vitro* and based on highly promising phase 2 data, with 98% cure rate, a global phase 3 program is ongoing as a combination of BEM and the HCV NS5 inhibitor ruzasvir^17^. These results prompted us to further validate its efficacy against HEV.

First, we validated that BEM inhibits replication of HEV-3 ORF2::GFP11 (green line) in a dose-dependent manner without impairing cell viability (gray line, **Figure 2C**). We confirmed the inhibitory capacity of BEM on the original cell-culture adapted HEV-3 Kernow C1 p6 strain (HEV-cc) by infecting S10-3 cells and determined a half-maximal inhibitory concentration (IC_50_) of 0.54 μM (**Figure 2D**). Finally, we found that the treatment decreased the release of HEV RNA into the cell culture supernatant of HEVcc-infected cells (**Supplementary Figure 2**).

Previously, we demonstrated that human-induced pluripotent stem cell-derived hepatocyte-like cells (HLCs) are permissive to authentic and non-adapted HEV isolate infection^26^. Using HLCs, we confirmed that BEM inhibited, to an even greater degree than NITD008, the replication of both, HEVcc (**Figure 2E**) and an HEV-3 isolate obtained from a patient’s stool (**Figure 2F**).

To determine the genetic barrier to resistance of BEM, we exposed HEVcc-infected S10-3 cells to varying concentrations of BEM every three passages for a total of 40 passages over 160 days (**Figure 2G**). Throughout this period, HEVcc replication remained sensitive to BEM, as evidenced by a consistent decrease in viral genome copies at either dose, even during the final three passages.

### Combining bemnifosbuvir with ribavirin has an additive inhibitory effect on HEV RNA replication in primary human hepatocytes

Next, we wanted to validate the potency of BEM to inhibit HEV replication in primary human hepatocytes (PHHs). In addition, we sought to analyze the combination of BEM with ribavirin, which is given to chronic HEV patients in the clinic.

Treatment of HEVcc-infected PHHs with BEM showed a dose-dependent inhibition of HEV replication as evidenced by a decrease of HEV genomes in the cell culture supernatant (**Figure 3A**). Immunofluorescence staining also showed reduction of ORF2 positive cells after BEM treatment (**Figure 3B, C**). Notably, BEM showed no cytotoxicity up to 10 μM in PHHs (**Supplementary figure 3**).

**Figure 3.**
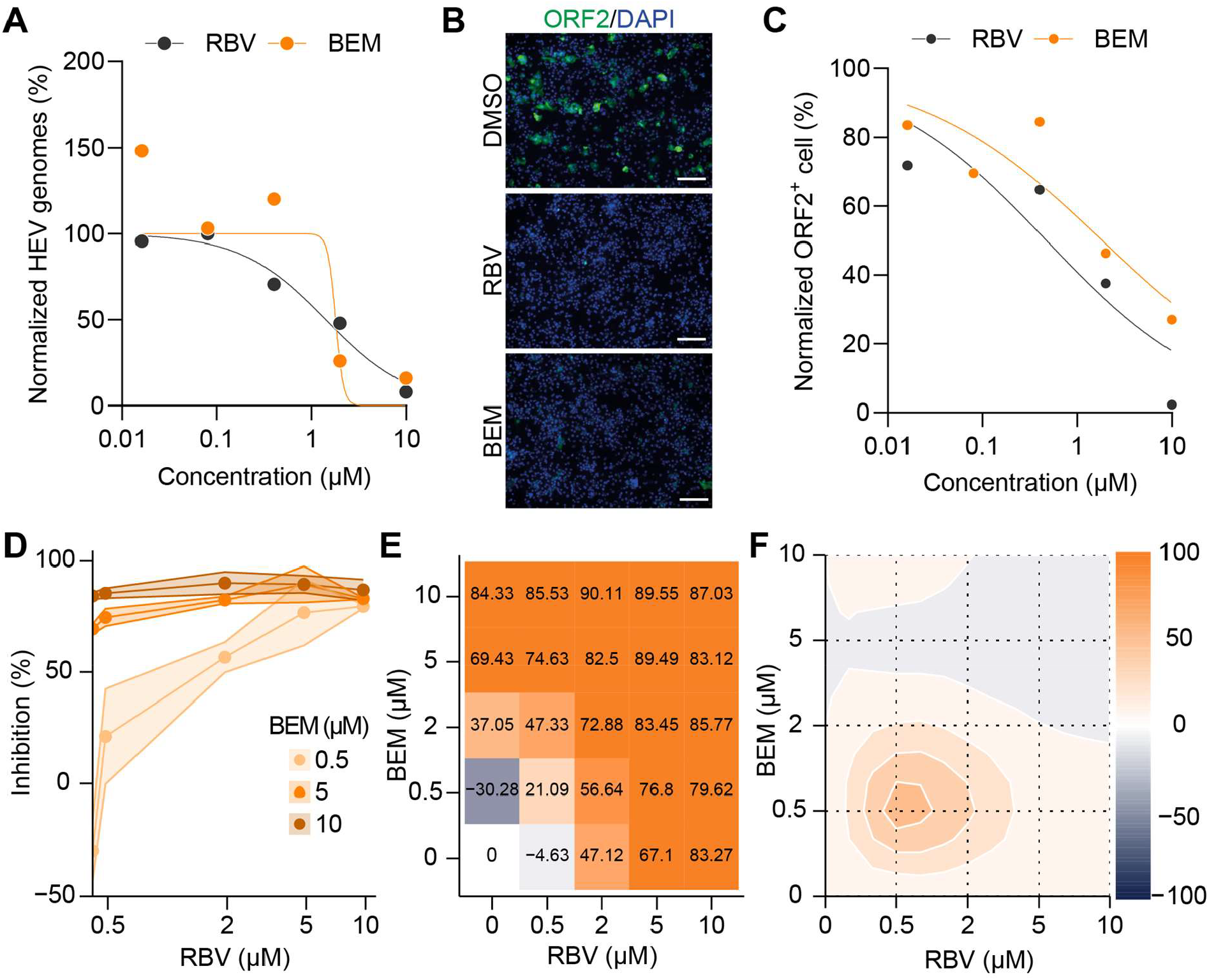
Combining bemnifosbuvir with ribavirin has an additive inhibitory effect on HEV RNA replication in primary human hepatocytes. (**A-C**), Primary human hepatocytes (PHHs) were infected with HEV-3cc WT (MOI = 0.05 FFU/well) and simultaneously treated with increasing concentrations of BEM for 72 h. DMSO was used as vehicle control. n = 1 donor. (**A**) PHH supernatants were harvested and extracellular HEV genomes were quantified by RT-qPCR. (**B-C**) HEV-3-infected and BEM-treated PHHs were fixed, stained against ORF2, and imaged on a widefield Keyence microscope. (**B**) Shown are representative images of PHHs stained for DAPI (blue, nucleus) and HEV ORF2 (green). Scale bar = 200 µm. (**C**) ORF2 positive cells were counted using the CellProfiler software. n = 20 microscope fields. (**D-F**), PHHs were infected with HEV-3cc WT (MOI=0.05 FFU/well) and treated with indicated molar concentrations of BEM, RBV, or the combinations of both. n = 2 donors. (**D**), HEV inhibition during treatment with 0.5 µM, 5 µM or 10 µM BEM and simultaneous titration of RBV (from 0 to 10 µM). (**E**), The dose-response matrix for RBV + BEM treatment depicted as the percent inhibition of viral infection. (**F**), A two-dimensional map of synergy scores depicting the combination of BEM with RBV.

With a mean zero interaction potency (ZIP) synergy score of 3.56 (where < −10 indicates antagonistic effects, −10 to 10 indicates additive effects, and > 10 indicates synergetic effects), BEM demonstrated additive antiviral effects when combined with RBV (**Figure 3D-F**). However, we also observed moderate synergistic effects when cells were treated with low doses (0.5 µM) of both BEM and RBV (**Figure 3F**).

### Bemnifosbuvir reduces HEV load and liver inflammation in a gerbil infection model

Finally, our goal was to evaluate the efficacy of BEM *in vivo* by determining viral loads upon BEM treatment in a gerbil HEV infection model^19^ (**Figure 4A**). The gerbils were inoculated with an HEV-3 isolate at 1×10^7^ genome copies per gerbil and were randomly divided into two groups (n = 3 per group). The treatment group received BEM at 250 mg/kg/day for 10 days, starting on day 5 post-inoculation. The vehicle treatment group received the same volume of solvent.

**Figure 4.**
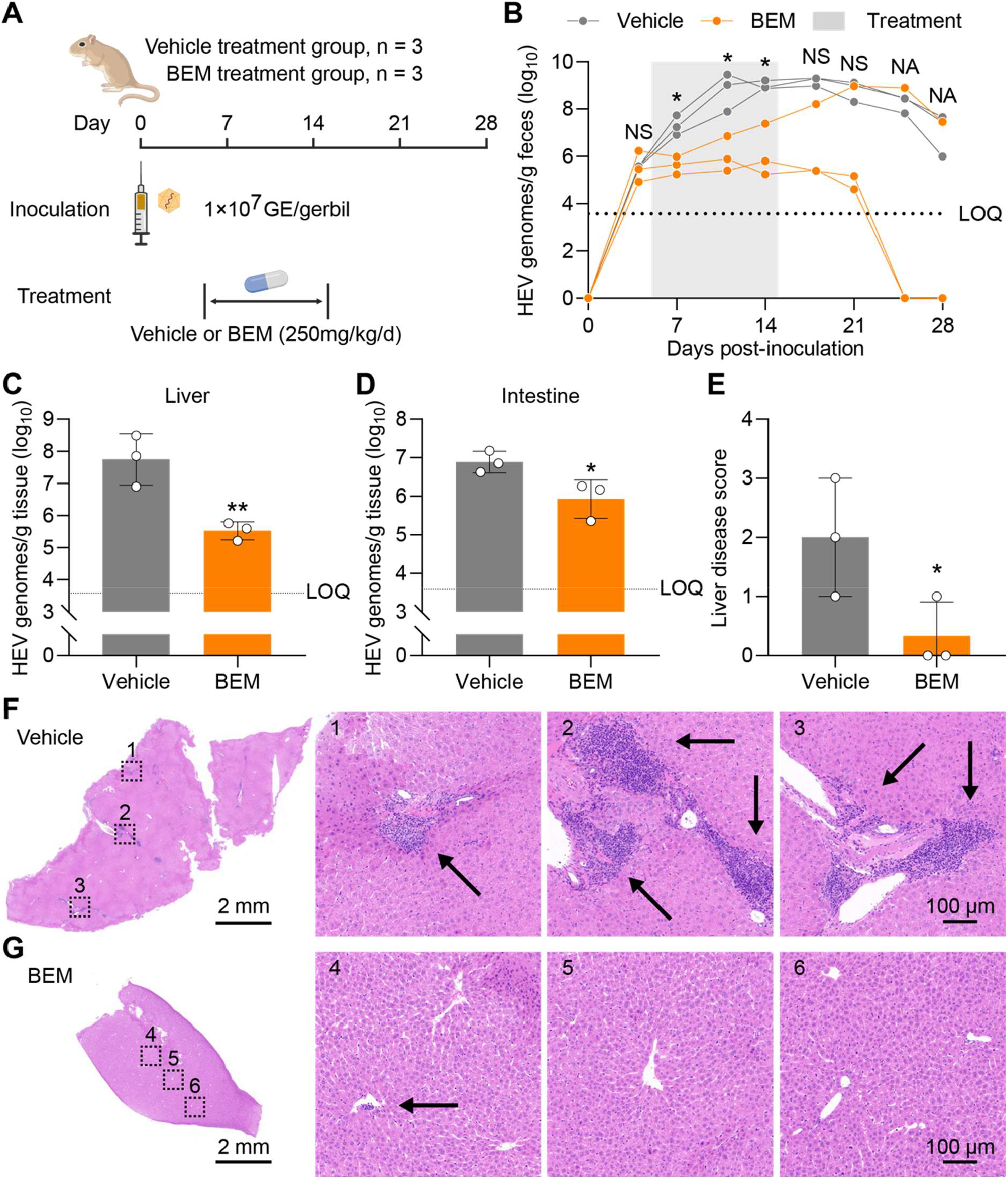
Bemnifosbuvir reduces HEV viral load and liver inflammation in a gerbil infection model. (**A**) Schematic illustration of the experimental design. Created in BioRender. Dao Thi, V. (2025) https://BioRender.com/aw6qce8 (**B**) HEV genomes were analyzed by RT-qPCR in feces samples collected at the indicated day post-infection. Statistical analysis was performed using unpaired Student’s t-test of each time point independently, adjusted for multiple testing using the Benjamini-Hochberg method. *, p < 0.05; **, p < 0.01; NS, non-significant. NA, non-applicable. (**C-D**) The liver (C) and (D) intestinal tissue samples were collected after euthaniasia and the HEV genomes were analyzed by RT-qPCR. Statistical analysis was performed using unpaired one-tailed Student’s t-test. *, p < 0.05; **, p < 0.01. (**E**) Degree of liver damage as assessed by Ishak scoring to analyze the H&E staining images of the liver tissue slices. Statistical analysis was performed using unpaired one-tailed Student’s t-test as we hypothesized the BEM treatment would decrease the Ishak scores. *, p < 0.05. (**F-G**) Shown are representative images of H&E stained liver tissue sections from vehicle treatment group (F) or BEM treatment (G) group, imaged with a standard light microscope. Black arrows, inflammatory infiltrates. Scale bar sizes are indicated in the images.

Throughout the treatment period, the fecal samples of all three gerbils in the BEM treatment group had significantly lower HEV RNA levels than those in the vehicle-treated group (**Figure 4B**). After discontinuing the treatment, one gerbil experienced a rebound, shedding HEV RNA at levels similar to those of the vehicle-treated gerbils from days 21 through 28 post-inoculation. In contrast, two gerbils achieved complete viral clearance more rapidly than those in the vehicle group and exhibited no detectable HEV RNA shedding by day 24 post-infection, despite the discontinuation of treatment.

The gerbils were euthanized on day 28 post-inoculation and we determined viral loads in the liver and intestinal tissues. As shown in **Figure 4C-D**, the viral loads were significantly lower in the BEM-treated group compared to the vehicle-treated group in both tissues. We then performed hematoxylin and eosin (H&E) staining on liver tissues from both groups of gerbils and analyzed them using the Ishak scoring system^27^. Animals in the BEM-treated group had significantly lower Ishak scores than those in the vehicle-treated group (**Figure 4E**). We found inflammatory infiltrates in the vehicle treatment group’s liver tissues, while we only occasionally found mild inflammatory infiltrates in the BEM treatment group (**Figure 4F-G** and **Supplementary Figure 4A-C)**. Of note, the gerbil with viral rebound showed moderate inflammatory infiltrates in the liver tissue (**Supplementary Figure 4D**). Altogether, these results demonstrated that BEM decreased HEV-induced liver inflammation and exhibited a protective effect.

## Discussion

The urgent need for effective and curative treatments for patients with chronic HEV infection remains a significant challenge. In this study, we developed a rapid and convenient high-content image-based screening platform utilizing a novel, full-length HEV-3 fluorescence reporter virus. This platform eliminates the need for biochemical assays or antibody-based staining of viral proteins. Furthermore, concurrently assessing antiviral activity (by measuring the inhibition of the GFP signal) and cytotoxicity (by quantifying cells using Hoechst staining) eliminates the need for separate assays.

Although we did not employ a suitable structurally diverse compound library in this initial work, future studies should leverage this platform to identify molecules that target HEV proteins ORF2 and ORF3. A prominent example is hepatitis B virus (HBV), where recent development of antiviral agents targeting the capsid protein has proven effective; these inhibitors disrupt capsid assembly, thereby preventing viral replication and infection^28^. Given the critical role of ORF2 in antagonizing host cell-intrinsic antiviral responses^15^, the HEV capsid ORF2 emerges as a highly compelling target for antiviral intervention.

In this study, we utilized our image-based screening platform to evaluate a library of nucleotide/nucleoside analogs and identified BEM as a potent inhibitor of HEV replication. BEM, previously characterized as a nucleotide analog with activity against other RNA viruses, exhibited dose-dependent suppression of HEV replication *in vitro* and *in vivo*, with minimal cytotoxicity at effective concentrations. The identification of BEM as an HEV inhibitor therefore expands the repertoire of candidate compounds and highlights the potential for repurposing existing antivirals.

BEM is a 2’-fluoro-2’-methyl modified guanosine nucleotide prodrug that is metabolized into its active 2’-fluoro-2’-methyl-GTP form^29^. In its active form, BEM can interfere with viral RNA synthesis by acting as an immediate chain terminator at the RdRp active site^30^. Importantly, studies have demonstrated that BEM is well tolerated in animal models^18^. Additionally, a Phase I clinical trial showed that BEM doses of up to 550 mg/day for seven days had minimal adverse effects^31^.

Compared to other nucleoside/nucleotide analog antivirals, BEM demonstrated greater activity against HEV in S10-3 hepatoma cells. BEM has an IC50 of 0.54 μM, which appears more potent than SOF (IC50 = 1.2 μM) and RBV (IC50 = 8 μM)^11^. Although NITD008 exhibits higher potency with an IC50 of 0.03 μM^24^, it has been associated with toxicity in animal studies ^25^.

In addition, combining BEM with RBV resulted in an additive antiviral effect against HEV. When both drugs were applied at low doses, we even observed a moderate synergistic effect, suggesting that combination therapy may enhance treatment efficacy. Finally, the passaging experiment indicated that HEV-3 maintains susceptibility to BEM over prolonged periods, reducing concerns about rapid resistance development as previously described for SOF and RBV^8-10 32^.

In the gerbil infection model, BEM reduced fecal viral loads during the treatment period and resulted in faster viral clearance. However, one gerbil experienced a viral rebound after treatment ended, suggesting the regimen needs optimization. This could include adjusting the dosage, schedule, and duration of treatment, or combining it with RBV. We acknowledge that the used gerbil model represents a model of acute HEV infection. Further investigations using chronic animal infection models are therefore warranted to evaluate the efficacy of BEM monotherapy or combination therapy in clearing chronic HEV infection. Nevertheless, BEM reduced liver inflammation and showed protective effects in the current model. Owing to BEM’s favorable safety profile in preclinical and clinical settings, our data therefore suggest the need for further investigation into its efficacy in treating HEV-infected patients.

In conclusion, our study introduces a versatile HEV reporter system and identifies BEM as a promising candidate for HEV therapy. These findings pave the way for further preclinical development and underscore the utility of innovative molecular tools in antiviral drug discovery.

## Acknowledgements

We thank Suzanne Emerson, Hans-Georg Kräusslich, Rainer G. Ulrich, and Stephen Duncan for sharing reagents. We thank Yi Zhang for sharing his expertise on the histological analysis. We acknowledge microscopy support from the Infectious Diseases Imaging Platform (IDIP) at the Center for Integrative Infectious Disease Research, Heidelberg, Germany and support from Daniel Kirrmaier and Michael Knop at the ZMBH, Heidelberg, Germany.

## Authors contributions

Conceptualization, J.H., V.L.D.T.; Methodology, J.H., T.L., M.K., E.S., L.W., V.L.D.T.; Investigation, J.H., T.L., M.K., A.F., E.T., H.C., P.J., X.Y., J.B.; Resources, V.L.; Formal analysis, J.H., T.L., M.K., H.C., E.S., L.W., V.L.D.T.; Writing – original draft preparation, J.H., V.L.D.T.; Final draft, J.H., V.L.D.T.; Supervision, E.S., L.W., V.L.D.T. Funding acquisition, E.S., L.W., V.L.D.T.

