## Supplementary for "The nucleotide analog bemnifosbuvir inhibits hepatitis E virus replication in preclinical models"

### Supplementary Methods & Materials

*Cell culture*

The human hepatoma cell line S10-3 (a kind gift from Suzanne Emerson) and HepG2/C3A (ATCC HB-8065) were grown in Dulbecco's Modified Eagle Medium (DMEM) (Gibco), supplemented with 10% fetal bovine serum (FBS) (Capricorn Scientific), referred to as complete DMEM (cDMEM). Collagen-coated cell culture plasticware was used for HepG2/C3A cells. The cells were cultured at 37 °C in a humidified incubator with 5% $CO_2$. S10-3 GFP1-10 cells were generated by lentiviral transduction based on a pWPI plasmid encoding GFP1-10. The cells were selected with 400 µg/mL G418 and continuously cultured under selection pressure.

The induced pluripotent stem cell line iPS.3CA (a kind gift from Stephen Duncan) was differentiated into hepatocyte-like cells (HLCs) as previously reported[1]. Briefly, iPSCs were seeded into Matrigel-coated culture plates for definitive endoderm (DE) differentiation using the STEMdiff Definitive Endoderm Differentiation Kit (Stem Cell Technologies). DE cells were seeded into Matrigel-coated culture plates for HLCs differentiation: 5 days in basal Rosewell Park Memorial Institute (RPMI) 1640 medium with HEPES (Gibco), supplemented with B-27 (Gibco), GlutaMAX (Gibco), NEAA (Gibco), pen/strep (Gibco), bone morphogenetic protein 4 (PeproTech) and fibroblast growth factor basic (Gibco), then 5 days in basal RPMI 1640 medium containing human hepatocyte growth factor (PeproTech), and finally 5 days in Hepatocyte Culture Medium BulletKit (Lonza) supplemented with human oncostatin M (R&D Systems).

Primary human hepatocytes (PHHs) were purchased from Primacyt (Schwerin, Germany) as cryopreserved hepatocytes and thawed according to the manufacturer's instructions. PHHs were seeded on 24-well plates and kept in Human Hepatocyte Maintenance Medium (HHMM, Primacyt). Donors were serologically tested and found negative for HIV, hepatitis B and C, and SARS-CoV-2.

*In vitro transcription (IVT) of HEV RNA genome and electroporation*

HEV plasmids were linearized with MluI (New England Biolabs) and RNA was *in vitro* transcribed using the mMESSAGE mMACHINE T7 kit (Invitrogen). $4\times10^6$ S10-3 or S10-3 GFP1-10 cells were electroporated with 1 µg (to analyze genome replication) or 10 µg (to produce virus and analyze drug inhibition) IVT HEV RNA in 400 µL cytomix (120 mM KCl, 0.15 mM $CaCl_2$, 10 mM $KPO_4$, 25 mM HEPES, 2 mM EGTA, and 5 mM $MgCl_2$) freshly supplemented with 2 mM adenosine triphosphate (ATP) and 5 mM glutathione (GT) using a Gene Pulser Xcell Electroporation System (Bio-Rad Laboratories). After electroporation, the cells were immediately transferred to pre-warmed cDMEM, and plated in cell culture flasks or plates.

*HEV virus production/isolation*

Virus from cells electroporated with HEV IVT RNA was harvested 7 days post-electroporation either from the cell lysate (intracellular naked HEV) or from the cell culture supernatant (extracellular quasi-enveloped HEV). Naked HEV particles were harvested from lysates by freeze-thaw cycles and concentrated by ultracentrifugation at 100,000 x g using a SW 32 Ti Swinging-Bucket Rotor for 3 h at 4 °C through a 20% sucrose cushion. The pellet was resuspended in PBS, aliquoted and stored at -80 °C. For quasi-enveloped HEV, cell culture supernatants were centrifuged at 9,000 g for 5 min to remove cell debris, aliquoted and stored at -80 °C.

Patient-derived HEV isolates for HLC and gerbil infection (GenBank accession number OM780137.1) were isolated from stool samples by resuspension in PBS and multiple centrifuge steps. The clarified supernatant was filtered through 0.45 and 0.22 µm filters, aliquoted, and stored at -80 °C.

*Virus titration*

Foci forming unit (FFU): HepG2/C3A cells were seeded in 48-well plates as $3\times10^4$ cells/well. On the next day, the cells were inoculated with virus in MEM medium (Gibco) with 10% FBS. The inoculum was removed 24 h later, and the cells were maintained in DMEM medium with 10% FBS. The cells were fixed with 4% paraformaldehyde (PFA) on day 5 after infection. Immunofluorescence staining against ORF2 protein was performed to visualize the infected cells. The cells were imaged with a CellDiscoverer 7 automated microscope (Zeiss). Foci were counted manually, and the virus titer was calculated.

Genome copy: HEV RNA genomes from the virus preps were extracted using Trizol according to the manufacturer's protocol. The genome copy number was quantified by RT-qPCR.

*RNA extraction and quantitative reverse-transcription PCR (qRT-PCR)*

Total RNA from cell lysates was isolated using either the Universal RNA Kit (Roboklon) or Trizol (Invitrogen™, Thermo Fisher Scientific), and RNA from the cell culture supernatant was isolated using Trizol, according to the manufacturer's protocol respectively. The RNA was reverse transcribed into cDNA using an iScript™ cDNA Synthesis Kit (Bio-Rad), and HEV genomes and the RSP11 housekeeping gene for normalization were quantified by qPCR using an iTaq Universal SYBR Green Supermix (Bio-Rad) on a CFX96 Touch Real-Time PCR Detection System (Bio-Rad) with primers listed in Supplementary table 1.

Total RNA from feces samples or from tissue samples were isolated using the EasyPure viral RNA/DNA kit (TransGen Biotech, Beijing, China). HEV viral load was determined using a one-step real-time quantitative PCR assay (GoTaq® Probe 1-step RT-qPCR System Kit, Promega, Wisconsin, USA) with primers listed in Supplementary table 1.

*Western blotting*

The cell culture supernatant from electroporated cells S10-3 GFP1-10 cells were collected, and cleared by centrifugation at 300× g for 5 min at 4 °C. The cells were lysed in Pierce RIPA buffer (Thermo Fisher Scientific) with cOmplete™ Mini Protease Inhibitor Cocktail (Roche) on ice for 30 min, followed by centrifugation at 17,000× g for 15 min at 4 °C. The cell culture supernatant and cell lysate samples were supplemented with Laemmli SDS sample buffer and boiled at 95 °C for 10 min.

Samples were run on SDS-polyacrylamide gel electrophoresis (PAGE) gels (5% stacking gel and 10% resolving gel). Proteins were transferred to PVDF membranes. The membranes were blocked with 5% milk (Carl Roth) in PBS/0.1% Tween-20 (PBS-T), and incubated with primary antibodies diluted in PBS-T overnight at 4 °C. After three washes with PBS-T, membranes were incubated with horseradish peroxidase (HRP)-coupled anti-mouse or anti-rabbit secondary antibodies (Jackson Immunoresearch, 1:4000) for 1 h at room temperature. After three washed with PBS-T and one wash with PBS, the chemiluminescent signals were developed using Pierce™ ECL Western Blotting Substrate (Thermo Fisher Scientific) or SuperSignal™ West Femto Maximum Sensitivity substrate (Thermo Fisher Scientific).

*Immunofluorescence staining and imaging*

Cells were fixed with 4% PFA for 10 min at RT, followed by permeabilization and blocking in PBTG (0.1% Triton X-100, 1% bovine serum albumin, and 10% goat serum in PBS) for 1 h at RT. Then, the cells were incubated with the primary antibodies in PBTG overnight at 4 °C. After washing with PBS, the cells were incubated with an Alexa Fluor™-conjugated secondary antibody (Thermo Fisher Scientific), diluted in PBTG, for 1 h at RT. The nuclei were stained with Hoechst 33342 (Thermo Fisher Scientific). Antibodies are listed in Supplementary table 2.

The cells were imaged with a LSM900 confocal microscope with an Airyscan 2 detector (Zeiss) or with a wide-field fluorescence microscope (Keyence BZ-X800E). The percentage of ORF2 positive ($ORF2^+$) cells was analyzed using Cellprofiler software.

*Cell viability assay*

$6\times10^3$ S10-3 cells were seeded into a well of a 96-well plate. The next day, the drugs and DMSO (as a control) were added to the cells. Three days later, CellTiter 96® AQueous One Solution (Promega) was added. After 1.5 hours, the absorbance at 490 nm was measured using a Tecan Infinite® M200 PRO plate reader. Cell viability was normalized to the DMSO control.

*LDH release assay*

Cell viability of PHHs was determined using the CytoTox 96® Non-Radioactive Cytotoxicity Assay (Promega). Lactate dehydrogenase (LDH) release in the supernatant was determined 72 h post infection by transferring 50 µL supernatant to a 96-well plate, followed by the addition of an equal volume of CytoTox 96® Reagent to each well. The reaction was incubated for 30 min and stopped by the addition of Stop Solution. Absorbance signal was measured at 492 nm in a plate reader (Tecan).

*Histology*

Tissues were fixed in 10% neutral buffered formalin for 3 days, embedded in paraffin, and cut into 5 µm serial sections. Hematoxylin and eosin (H&E) staining was performed. The samples were photographed and analyzed under a CX31 light microscope (Olympus) equipped with a digital camera with the assistance of CaseViewer software.

*Declaration of generative AI and AI-assisted technologies in the writing process*

During the preparation of this work, the authors used DeepL Write in order to improve language and readability. After using this tool/service, the authors reviewed and edited the content as needed and take full responsibility for the content of the publication.

**Supplementary Tables**

**Supplementary table 1: Primers used in this study**

| Name | Purpose/Target | Sequence (5’→3’) | Reference |
|---|---|---|---|
| ORF2 GFP11 fw | Cloning / ORF2::GFP11 | GACCACATGGTCCTTCATGA GTATGTAAATGCTGCTGGGA TTACTTAATGACAGTCCACTA TTGCTG | This study |
| ORF2 GFP11 rv | | GAAGGACCATGTGGTCACG ACTTCCACCACCGCCTGAAC CTCCACCGCCAGACTCCCG GGTTTTG | |
| ORF2 SnaBI fw | | GTATGGGTCGTCTACCAACC | |
| ORF2 BbsI rv | | AAAAGGAAGACTGCGAAGG GGGCACG | |
| HEV fw | qPCR / HEV | GGTGGTTTCTGGGGTGAC | [1] |
| HEV rv | | AGGGGTTGGTTGGATGAA | |
| HEV probe | | TGATTCTCAGCCCTTCGC | [2] |
| RPS11 fw | qPCR / RPS11 | GCCGAGACTATCTGCACTAC | [1] |
| RPS11 rv | | ATGTCCAGCCTCAGAACTTC | |

**Supplementary table 2: Antibodies used in this study**

| Antibody | Application & dilution | Source |
|---|---|---|
| Mouse anti-ORF2 | IF, 1:400<br>WB, 1:1000 | MAB8002, Merck |
| Rabbit anti-ORF2 | IF, 1:5000 | A kind gift from Rainer G. Ulrich, Friedrich Loeffler Institute, Greifswald, Germany |
| Rabbit anti-GFP11 | WB, 1:1000 | PA5-109258, Invitrogen |
| Mouse anti-ORF3 | WB, 1:25 | ABCD_RB198, University of Geneva Antibody Facility |
| Mouse anti-β-Actin | WB, 1:4000 | A2228, Sigma-Aldrich |
| Rabbit anti-GFP | IF, 1:2500 | A kind gift from Hans-Georg Kräusslich, CIID, Heidelberg, Germany |

**Supplementary figures and figure legends**

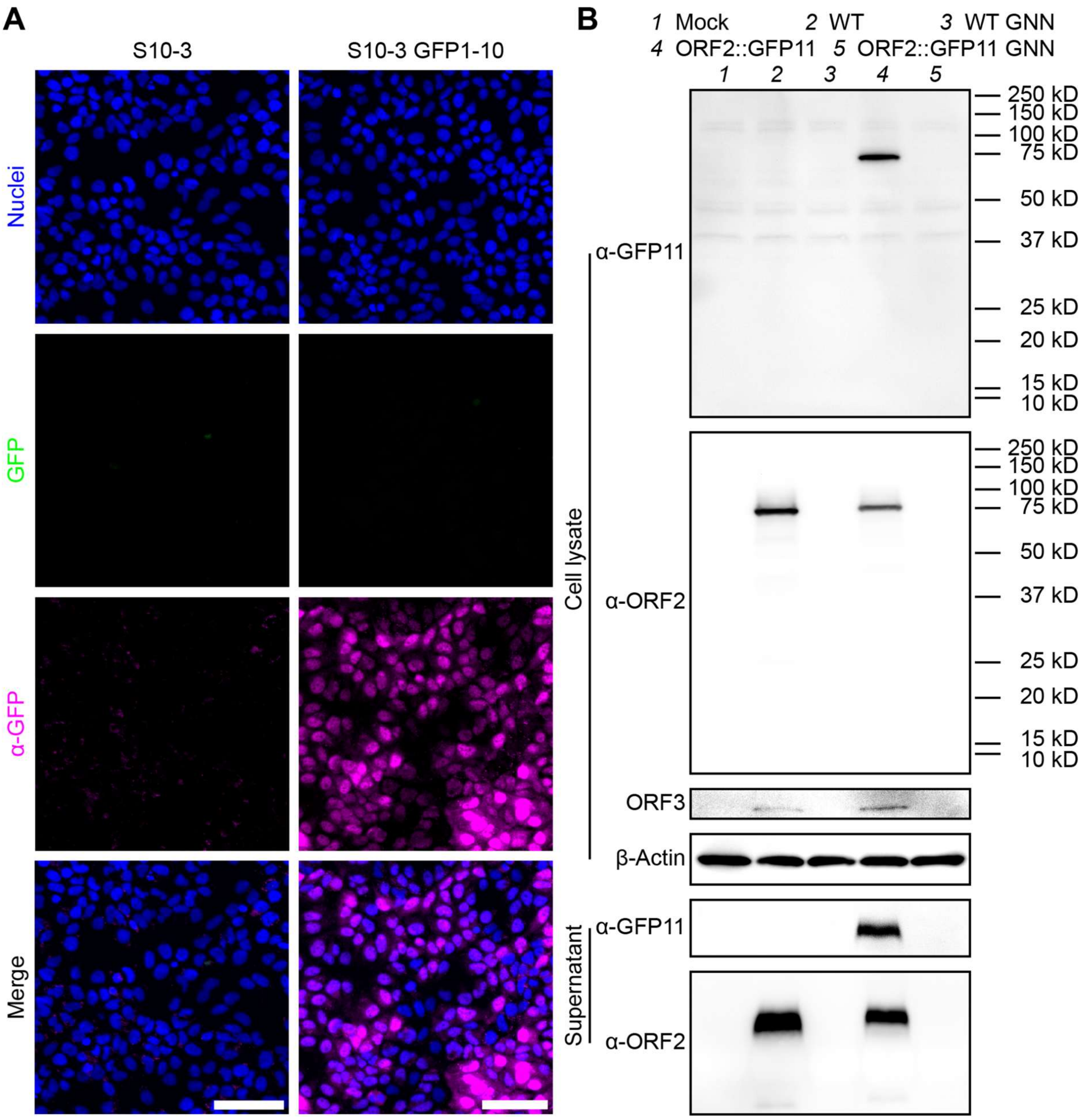


**Supplementary figure 1. S10-3 GFP1-10 cells express GFP1-10 protein; HEV-3 ORF2::GFP11 expresses ORF2::GFP11 and ORF3 proteins.** (**A**) S10-3 and S10-3 GFP1-10 cells were fixed, stained for nucleus (blue) and GFP1-10 (magenta), and imaged on a widefield CellDiscoverer7 microscope. Shown are representative images. Scale bar = 100 µm. (**B**) S10-3 GFP1-10 cells were electroporated with *in vitro* transcribed HEV RNA. Western blot analysis for β-Actin, ORF2, and ORF3 in the cell lysate, and ORF2 in the cell culture supernatant were performed.

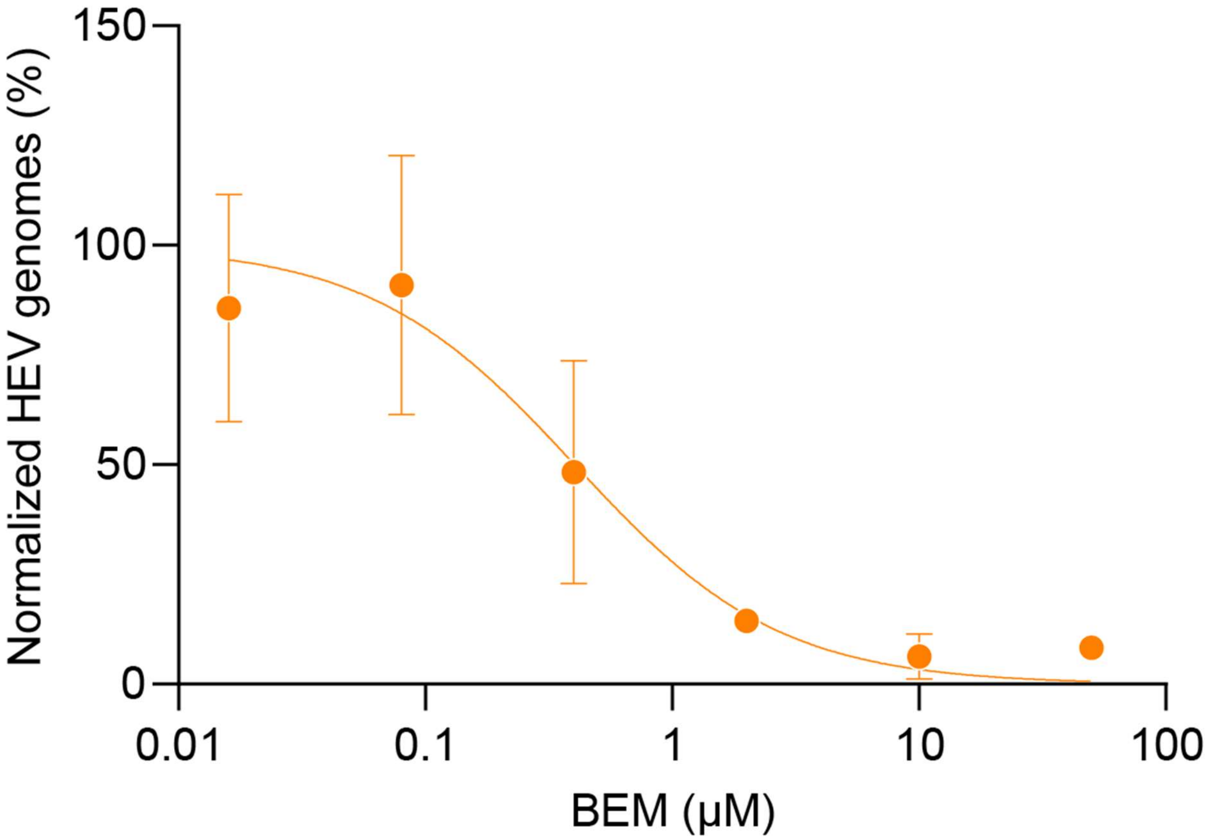


**Supplementary figure 2. BEM decreases HEV RNA in the cell culture supernatant in a dose-dependent manner**. S10-3 cells were infected with HEV-3cc WT virus (MOI=1 FFU/cell). The next day, viral inoculum was removed and cells were treated with different concentrations of BEM for 72 h. Extracellular HEV genomes in the cell culture supernatant were isolated by Trizol, and quantified by RT-qPCR. 0.5% DMSO was used as vehicle control. Data show mean ± SD of n = biological replicates from 3 independent experiments.

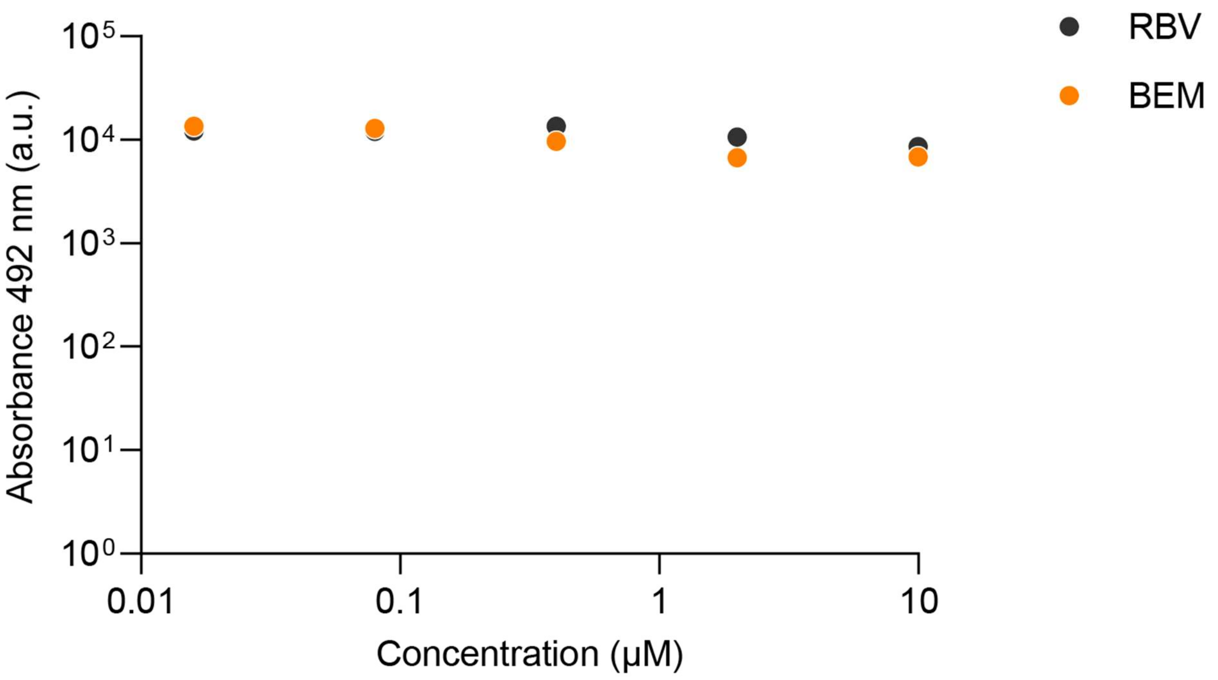


**Supplementary figure 3. BEM showed no cytotoxicity up to 10 μM in PHHs.** PHHs were seeded in a 24-well plate. The next day, cells were infected with HEV-3cc and incubated in the presence of BEM or RBV for 3 days. DMSO served as vehicle control. 50 μL supernatant was used for LDH release assay. n = 1 donor.

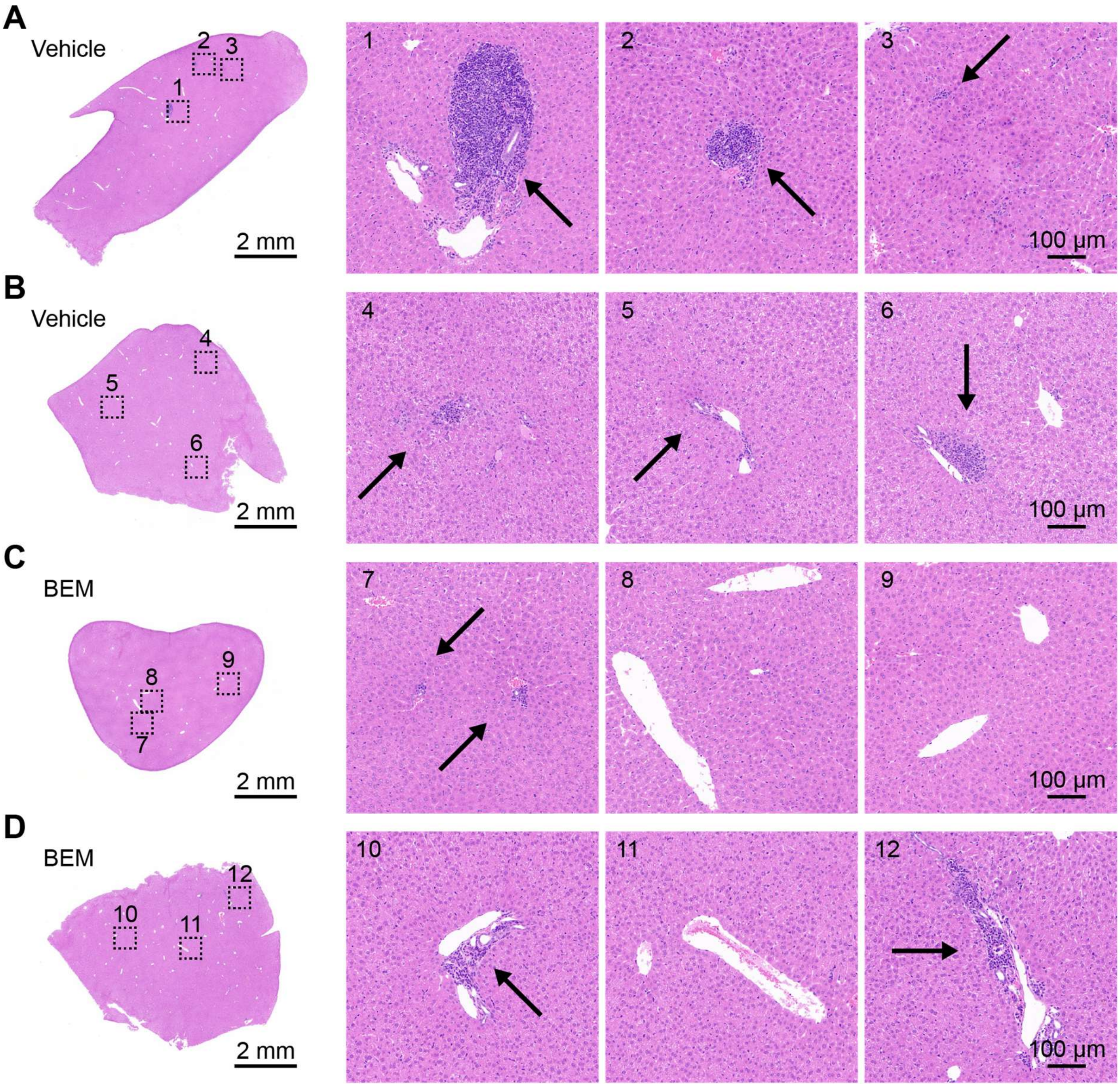


**Supplementary figure 4. BEM reduced liver inflammation in a gerbil infection model.** (A-D) Shown are the overview and zoom-in H&E staining images of the liver tissue slices from the other two gerbils in vehicle treatment (A-B) or BEM treatment (C-D) group. Black arrows, inflammatory infiltrates.
